# Automatic pain face analysis in mice: Applied to a varied dataset with non-standardized conditions

**DOI:** 10.64898/2026.02.16.706098

**Authors:** Niek Andresen, Manuel Wöllhaf, Jenny Wilzopolski, Annemarie Lang, Angelique Wolter, Laura Howe-Wittek, Clara Bekemeier, Leonie-Iva Pawlak, Sarah Beyer, Holger Cynis, Edith Hietel, Vera Rieckmann, Max Rieckmann, Christa Thöne-Reineke, Lars Lewejohann, Olaf Hellwich, Katharina Hohlbaum

**Author notes:** These authors contributed equally to this work (Niek Andresen, Manuel Wöllhaf).

## Abstract

Biomedical research relies on scientifically validated tools to assess pain, suffering, and distress in laboratory animals to ensure their well-being. In mice, the most frequently used laboratory animals, the Mouse Grimace Scale (MGS) provides a reliable tool for the assessment of facial expression changes caused by impaired well-being. However, no automated tool can yet reliably assess all features of the MGS across different mouse strains under varying experimental or housing conditions in real-time, as the variability present in recorded image datasets poses substantial challenges for computer vision models. Despite this technical difficulty, variability across subsets in terms of mouse strain, treatments, laboratory, and image acquisition setup is essential for paving the way toward MGS assessment under non-standardized conditions in the home cage rather than standardized cage-side recording setups. Against this background, a large and diverse dataset containing five subsets is introduced and a deep learning model was trained to predict the average MGS scores ranging between 0 and 2. It achieved a root mean squared error (RMSE) of 0.26 when trained on all subsets of the dataset, outperforming the average human rater in terms of error magnitude. The correlation between human raters and automated MGS scores was very high (Pearson’s r=0.85). In the cross-dataset evaluation, one subset was excluded from training and used for testing the model. This approach yielded higher errors compared to models trained and tested on the same subsets. A model restricted to the feature of orbital tightening showed lower performance than one trained on all facial features of the MGS. Overall, the most reliable model for predicting average MGS scores for a novel dataset is the one trained on the combined subsets. Performance may be further enhanced by fine-tuning the model using human-generated MGS scores for a portion of the novel subset.

## Introduction

Working with laboratory animals requires close monitoring of their well-being to meet both ethical and legal standards. In addition, the well-being of laboratory animals is essential for good scientific practice, as pain, suffering, and distress may substantially affect the data generated in an animal experiment and increase data variability [1, 2]. The ideal approach would be fully automated 24-hour monitoring of the animals’ well-being in their familiar environment, such as the home cage of laboratory mice. Automated 24-hour monitoring takes into account the circadian rhythm, i.e., nocturnal animals can be observed in their activity phase at night. When the behavior of nocturnal animals is assessed for a few hours during the light phase only, important information may be missed [3, 4]. Another advantage of automated systems is that the presence of humans is not required. It is assumed that in the presence of an experimenter, prey animals such as mice show altered behavior and mask, for instance, signs of pain [5]. Therefore, automated systems may be more sensitive to identify animals with a reduced well-being than live cage-side health checks operated by humans [6]. Moreover, automation enables a more objective and data-driven evaluation of the parameters and can reduce labor time [6].

Various behavioral, physiological, and external appearance-related parameters are investigated to assess the well-being of mice, the most commonly used laboratory animals [7, 8]. One of these parameters is facial expression, which can be analyzed using the Mouse Grimace Scale (MGS) [9]. The MGS consists of five facial action units (i.e., orbital tightening (OT), nose bulge, cheek bulge, ear position, and whisker change), which are assessed on a scale from 0 to 2 (0 = not present, 1 = moderately visible, 2 = severe). If a mouse is in pain, the facial action units shift toward higher scores. Severe pain results in closed eyelids and nose as well as cheek bulge [9]. In addition, the ears appear more pointed, rotate outward and/or backward, and the distance between them increases [9]. The whiskers lose their natural curvature and appear straight, and they may be drawn backward or pulled forward [9]. Currently, the MGS is predominantly applied manually by trained personnel. Consequently, the facial expression is assessed only over a short period of time and may be altered by the presence of a human observer. Full and reliable automation of the MGS application would allow for continuous 24-hour monitoring of mice’s facial expression to detect impairments in well-being at an early stage.

In recent years, various approaches have been developed towards automated MGS analysis [10–22]. The foundation was laid by Socotinal et al. who developed a face detection algorithm that can identify faces of white rats in images [10]. An automated picture selection tool for black-furred mice, such as C57BL/6J or C57BL/6N mice, developed by Ernst et al., enables the selection of high-quality images suitable for MGS assessment in a manner that is standardized and not influenced by the experimenter [11, 12]. The first step towards recognizing the well-being of mice from their facial expression was taken by Tuttle et al. through binary classification, which distinguished between mice with impaired and unimpaired well-being [13]. Tuttle et al. used images of white-furred mice (CD-1) [13]. They reached a validation accuracy of 94 % and also found a correlation between the classifier’s confidence output and the MGS score obtained by human raters [13]. Averaging the output over several images of the same moment boosts the accuracy of such a task, as shown by Andresen et al. for black-furred mice (C57BL/6J) [14]. With this approach, it was possible to achieve an accuracy of up to 99 % in a binary classification task with another mouse image dataset [14].

Kopaczka et al. [15] were the first to assign MGS scores for OT as a continuous value to faces of black-furred mice (C57BL/6). For this, a small dataset of 314 MGS-labeled images was used. They reported a mean absolute error of 0.871 on a finer grained 10-point scale ranging from 0 to 9 [15]. This corresponds to an error of 0.194 on the MGS scale with values from 0 to 2. In another study by Vidal et al., a network was trained on 1,087 MGS-labeled images of white-furred mice (ICR) to predict the score for OT on a 3-point scale from 0 to 2 [16]. The average accuracy was 69.3 % on image crops containing the eyes [16]. Gupta et al. also focused on OT, but chose a different approach by measuring eyelid distance as well as palpebral fissure width of black-furred mice (Nav1.8-ChR2) and providing a quantitative OT [17].

The next step was taken by Chiang et al. with the development of “DeepMGS” using white-furred mice (ICR, 1,504 MGS-labeled images), which automatically predicts MGS scores of all facial action units and computes the total sum [18]. “DeepMGS” achieved an accuracy between 70 and 90 %. While “DeepMGS” has not been made publicly available, the “PainFace” tool developed by McCoy et al. can be accessed freely online [19]. “PainFace” is based on a big dataset of more than 70,000 MGS scored images of black-furred mice (C57BL/6) [19]. It assesses OT, ear position, and whisker changes on a 3-point scale ranging from 0 to 2, and nose bulge on a binary scale, but regression to a continuous value was not successful. For the classification of feature scores, they do not report accuracy, but the F-score of at least 0.75. It is recommended to use a brightly lit observation chamber to apply “PainFace”, as described in detail in the study [19]. Recently, “PainFace” was used to develop an algorithm for white-furred mice (CD-1, *>* 63,000 MGS scored images) that detects all five facial action units on a 3-point scale from 0–2, with F-scores ranging from 0.67 to 0.84 [20].

Beyond retrospective MGS analysis of image data as described in the above-mentioned studies, Sturman et al. provide a real-time application called “GrimACE” [21]. The use of GrimACE requires a specific cage-side recording setup. It was created using a vision transformer network and trained on 1,245 MGS-labeled images of black-furred mice (BHiCre mice, C57BL/6 mice). The system includes a frame selector and a face detector. Performance is reported as a Pearson correlation of 0.87 between the model and a human expert rater. Besides MGS scores, “GrimACE” also provides pose estimation.

However, all automated tools for MGS assessment still require removing the mouse from its home cage and transferring it into a standardized cage-side recording setup, which causes additional distress and alters its behavior, including facial expression.Standardized cage-side recording setups are used because the variability in the recorded image data generated for MGS scoring is challenging for any computer vision model. Moreover, that is one of the the reason why, so far, no automatic tool can reliably assess each facial action unit of the MGS in different mouse strains under varying experimental or housing conditions in the home cage. Mice display diverse fur colors and are recorded in different cages under varying lighting conditions. The cages differ in dimensions, enrichment items, bedding, and nesting material resulting in a considerable variation in the background settings of the recordings. This leads to the problem that approaches of automated MGS systems may be limited to a mouse strain and certain experimental or housing conditions and cannot be transferred between laboratories. To create an automated system for facial expression analysis that can abstract from the mentioned parameters, an extensive dataset is needed. This dataset should include image material with a variety of mouse strains, procedures, and experimental or housing conditions.

In this work, we are addressing this issue and are using a large dataset (approximately 35,000 images) with non-standardized parameters: it contains MGS scores for images from five different mouse strains with albino, dilute brown, and black fur. The mice were subjected to different procedures in five different laboratories where various technical equipment for image acquisition (i.e., cameras, cage-side recording setup) was utilized. We applied current deep learning methods to automatically assign MGS scores, ranging from 0 to 2, to the images. The approach was based on transfer learning. A deep neural network was trained on pretext tasks first (i.e., the binary classification between mice with impaired and unimpaired well-being), and then acquired weights were used as initialization for the training of the main task. By making this large, diverse dataset and the deep learning model for automatic MGS scoring openly available, we aim to promote further efforts toward automated MGS assessment under non-standardized conditions, such as in the home cage.

## Materials and methods

This study presents a large dataset containing approximately 35,000 images of mouse faces. The dataset is publicly available under https://doi.org/10.14279/depositonce-25192 and is divided into five subsets, which were named by one of the authors involved: Table 1 summarizes the key information of the subsets and Fig 1 shows examples images of each subset. In the following, the individual subsets and the data processing steps are described, followed by details of the neural network model that was used.

**Table 1.** Overview of the five subsets.

|  | camera | mouse strain | sex | fur color |
| --- | --- | --- | --- | --- |
| AW | gray video | C57BL/6N | m | black |
| JW | gray video | BALB/c | f/m | albino |
| KH | RGB | C57BL/6J | f/m | black |
| LW | RGB | NMRI, C57BL/6J | f/m | albino, black |
| MR | RGB | DBA1 | m | dilute brown |
|  | frame selection | face detection | # images | # MGS labelled |
| AW | own method | DLC | 2,771 | 362 |
| JW | own method | DLC | 2,778 | 546 |
| KH | still photographs | DLC | 28,113 | 1,224 |
| LW | still photographs | manual | 597 | 597 |
| MR | still photographs | manual | 677 | 677 |

**Table 2.** Rater agreement.

(a) IRR
| subset | OT | NB | CB | WC | EP | avg |
| --- | --- | --- | --- | --- | --- | --- |
| AW | 0.40 | 0.38 | 0.31 | 0.34 | 0.24 | 0.33 |
| JW | 0.52 | 0.26 | 0.18 | 0.41 | 0.38 | 0.42 |
| KH | 0.52 | 0.15 | 0.14 | 0.30 | 0.31 | 0.28 |
| LW | 0.40 | 0.02 | 0.10 | 0.08 | 0.05 | 0.13 |
| MR | 0.73 | 0.22 | 0.18 | -0.07 | 0.35 | 0.28 |

(b) ICC
| subset | OT | NB | CB | WC | EP | avg |
| --- | --- | --- | --- | --- | --- | --- |
| AW | 0.37 | 0.61 | 0.42 | 0.43 | 0.26 | 0.42 |
| JW | 0.35 | 0.31 | 0.27 | 0.67 | 0.51 | 0.42 |
| KH | 0.53 | 0.20 | 0.19 | 0.24 | 0.25 | 0.28 |
| LW | 0.52 | 0.08 | 0.20 | 0.07 | 0.25 | 0.22 |
| MR | 0.88 | 0.22 | 0.29 | -0.12 | 0.23 | 0.30 |

(a) Inter-Rater Reliability (IRR) and (b) two-way mixed Intraclass Correlation Coefficient (ICC). OT: orbital tightening; NB: nose bulge; CB: cheek bulge; WC: whisker change; EP: ear position; avg: average.

**Fig 1.**
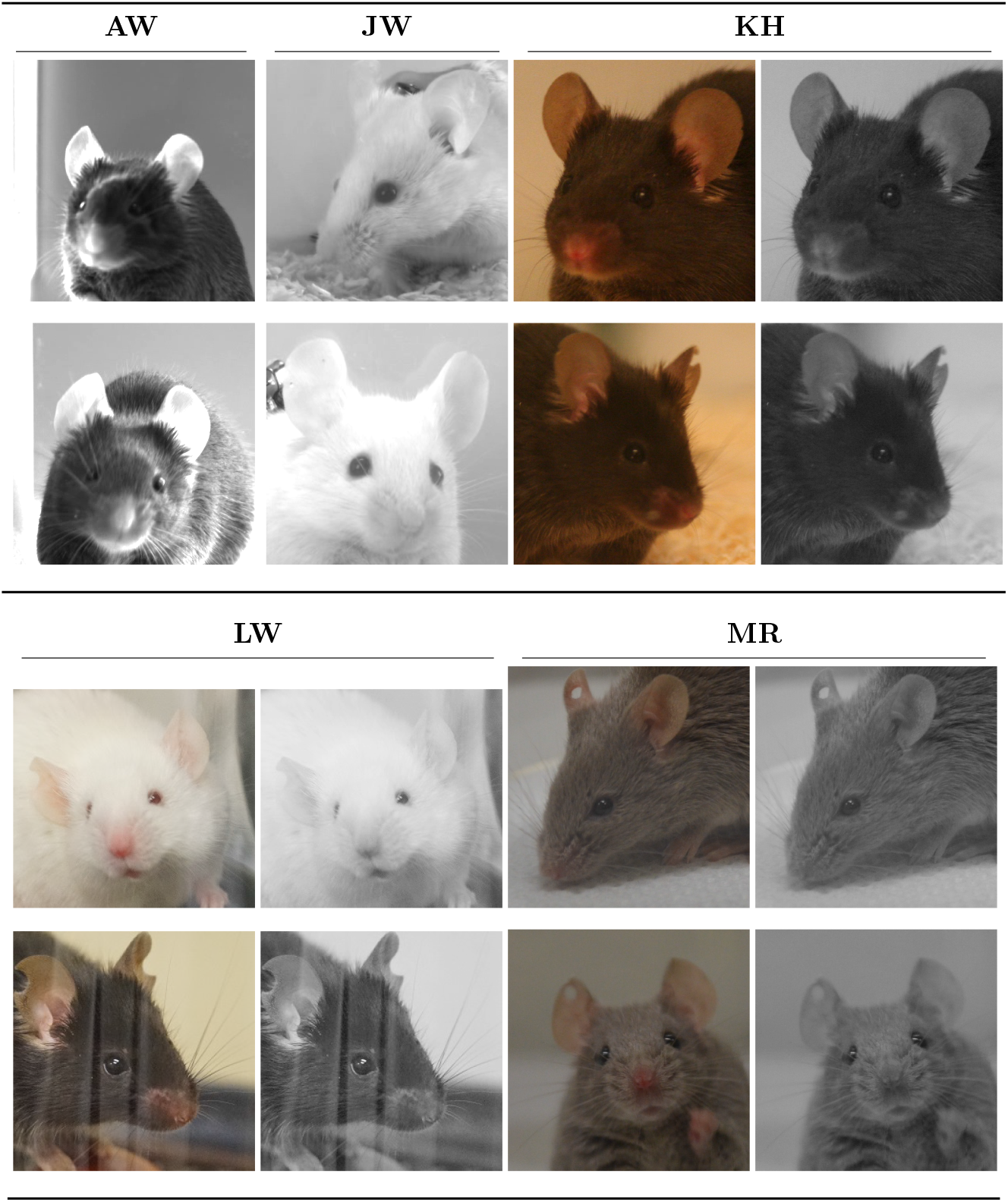
Examples from the five subsets. If the original image was acquired in color, the RGB image is shown alongside the corresponding grayscale version. The latter was used to train the neural network model. The subsets were named by one of the authors involved in image acquisition of the subset. AW: Angelique Wolter; JW: Jenny Wilzopolski; KH: Katharina Hohlbaum; LW: Laura Howe-Wittek; MR: Max Rieckmann.

“Own method” is described in the section on “frame selection and face detection”. DLC: DeepLabCut [23]; f: female; m: male; RGB: “Red, Green, Blue” color imagery.

### Subset AW

To generate the subset AW, pictures were taken from videos generated during a previously conducted and already published study [24]. Only videos of male mice with rigid and flexible fixation after femoral osteotomy/fracture receiving Tramadol analgesia in the drinking water were reused.

#### Animals

A total of 18 male C57BL/6N mice aged 10 weeks at the time of anesthesia/analgesia and 12 weeks at time of surgery were included in the study. Mice were either bred inhouse (Experimental Medicine Research Facilities; Charité - Universitätsmedizin Berlin, Berlin, Germany) or obtained from Charles River Laboratories (Sulzfeld, Germany). Health monitoring was regularly performed by the animal facility following the FELASA recommendations [25]. The animal experiment was approved by the Berlin state authority (Landesamt für Gesundheit und Soziales – LAGeSo; permit number: G0044/20) [24].

#### Housing and husbandry

Pairwise housing was conducted in IVC cages (Eurostandard Type II, Tecniplast, Buguggiate, Italy) in a semi-barrier facility with restricted entry through an airlock wearing personal protection cloth [24]. General housing was conducted with a 12:12 h light/dark cycle, a room temperature of 22 ± 2 °C, a humidity of 55 ± 10 %. Mice had free access to food (Standard mouse diet, Ssniff Spezialdiäten, Soest, Germany) and tap water [24]. A separated pair housing system was used (Green Line IVC Sealsafe PLUS Rat GR 900 cages, Tecniplast, Buguggiate, Italy) in case mice had to be separated due to aggressive behavior [24]. Cages were equipped with wooden chips (SAFE FS 14, Safe Bedding, Rosenberg, Germany), Envirodri (Shepherd Specialty Papers, USA), and shredded paper towel, a clear handling tube (Datesand Group, Bredbury, UK) and a double swing (Datesand Group, Bredbury, UK) [24]. Handling tunnels and double swings were removed after surgery to avoid potential injuries [24]. All experimenters handling the mice regularly for behavioral analysis were female and mice were habituated to the experimental setups and exclusively tunnel handled [24].

#### Treatments

Mice underwent two different interventions within two weeks. Baseline measurements were performed during training and habituation time. At 10 weeks of age, mice underwent anesthesia (15 min isoflurane inhalation anesthesia) and administration of analgesia without surgery/osteotomy to establish additional baseline measurements. At 12 weeks of age, osteotomy surgery was performed on the left femur using an either rigid or flexible external fixator for stabilization (rigid: 18.1 N/mm; flexible: 3.2 N/mm, both RISystem, Davos, Switzerland) [24]. The used surgical procedure has been described in detail previously ( [24], [26], [27]). For analgesia, mice received a subcutaneous injection of burprenorphine (1 mg/kg Temgesic, RB Pharmaceuticals, Heidelberg, Germany) before surgery and tramadol with the drinking water (0.1 mg/ml, Tramal Drops, Grünenthal, Stolberg, Germany) one day before and three days after surgery [24].

#### Image acquisition

Videos were taken at three baseline timepoints and 12 h, 24 h, 48 h, and 72 h post intervention (anesthesia/analgesia only; anesthesia/analgesia and osteotomy) using a high-resolution infrared camera (Basler Video Recording Software, Ahrensburg, Germany) [24]. For this, mice were transferred in clear observation boxes (10 cm x 10 cm x 14 cm; ground area = 100 cm^2^; modular animal enclosure; Ugo Basile, Gemonio, Italy) [24]. The clear observation boxes contained a soft pad as base that was changed regularly, and the inner surface of the observation boxes were cleaned after every mouse. Two mice were video taken next to each other in separate observation boxes. Mice were shortly habituated before videos were taken for 3 min [24]. Post-processing and analysis were carried out fully blinded.

### Subset JW

#### Animals

20 female and 20 male 5–6 weeks old BALB/c (BALB/cAnNCrl) mice were obtained from Charles River (Sulzfeld, Germany) [28]. Mice were acclimatized to their new housing environments for 14 days prior to the experiments [28]. The animals were free of all viral, bacterial, and parasitic pathogens listed in the FELASA recommendations [25]. Animal experiments were approved by the Berlin state authority (Landesamt für Gesundheit und Soziales – LAGeSo; permit number: G0204/20) [28].

#### Housing and husbandry

The mice were kept in a conventional facility at the Freie Universität Berlin. The mice were housed in groups of 3–4 animals in Makrolon type III cages (42 cm × 26 cm × 15 cm) with a wire lid. Wooden bedding material (SAFE® prime S, Rettenmaier & Söhne GmbH + Co. KG, Rosenberg, Germany), paper towels, and three cocoons (Datesand Group, Bredbury, UK) as nesting materials were provided. The cages were enriched with two red plastic houses (length: 100 mm, width: 90 mm, height: 55 mm; ZOONLAB GmbH, Castrop-Rauxel, Germany) and one metal tunnel (length: 125 mm, diameter: 50 mm). The mice were housed at a room temperature of 22 ± 2 °C and relative humidity of 55 ± 10 % on a light:dark cycle of 12:12 h (artificial light; lights on from 7:00 a.m. to 7:00 p.m.). A pelleted standard mouse diet (Altromin, Lage/Lippe, Germany) and tap water were available ad libitum. The animal technicians were both male and female. The experimenter was female. The mice were tail-handled. At the end of the experiments the mice were sacrificed for organ-/tissue-sampling. Mice were deeply anesthetized with ketamine-xylazine and exsanguinated via the V. cava caudalis.

#### Treatments

On day 0 of the experiment, Alzet micropumps 1007D (Charles River, Sulzfeld, Germany) were implanted under the back skin of the mice according to the protocol of Lu et al. [28, 29]. Mice were divided into 4 treatment groups in the first round (vehicle, treatment A, B & C; n = 3 mice/group & sex) for more details see Graff et al. 2024 [28]. In the second round the mice were divided into two groups (vehicle & treatment B; n = 4 mice/group & sex). On day 4 a blood sample (110 µl) was collected via the V. facialis [28].

#### Image acquisition

Image acquisition was performed with slight modifications according to Hohlbaum et al. [30]. The images were generated in a modified type-II-macrolon-cage (20,5 × 13 × 10,5 cm, two white walls, one transparent wall). The floor was covered with 0.5 cm fresh wooden bedding material. Before images were acquired, the mice were individually transferred into the observation cage. A Basler acA1920-40µm camera (Basler AG, Ahrensburg, Germany) was utilized for video acquisition. The videos were acquired via Basler video recording software 64-Bit (Basler AG, Ahrensburg, Germany) for approximately 1–2 minutes (in some cases more time was needed) at 41 frames/s with an exposure time of 9,800 µs. Mice were video monitored 30 min., 4 h, 12 h, and 24 h after the pump implantation. Prior to and directly after blood sampling via the V. facialis on day 4 and on day 7 before euthanasia (end of experiment).

### Subset KH

The subset KH is an extended version of the dataset previously described and used in Andresen et al. [14].

#### Animals

61 female and 65 male adult C57BL/6JRj mice were obtained from Janvier Labs (Saint-Berthevin Cedex, France) [31, 32]. The animals were free of all viral, bacterial, and parasitic pathogens listed in the FELASA recommendations [25]. Animal experimentation was approved by the Berlin state authority (Landesamt fü r Gesundheit und Soziales - LAGeSo; permit number: G0053/15).

#### Housing and husbandry

The mice were kept in a conventional facility at the Freie Universität Berlin. Females were housed in groups of 3–5 animals in Makrolon type IV cages (55 cm × 33 cm × 20 cm); males were individually housed in Makrolon type III cages (42 cm × 26 cm × 15 cm) because of aggressive behavior towards conspecifics [14, 31–33]. Fine wooden bedding material (LIGNOCEL® 3–4 S, J. Rettenmaier & Söhne GmbH + Co. KG, Rosenberg, Germany) and cotton nest material (nestlets: Ancare, UK agents, Lillico, United Kingdom) were used [14, 31, 32]. Additionally, cocoons were provided for castrated mice (ZOONLAB GmbH, Castrop-Rauxel, Germany) [14, 33]. The cages were enriched with a red plastic house (length: 100 mm, width: 90 mm, height: 55 mm; ZOONLAB GmbH, Castrop-Rauxel, Germany) and metal tunnels (length: 125 mm, diameter: 50 mm; one tunnel in Makrolon type III cages, two tunnels in Makrolon type IV cages) [14, 31–33]. The mice were housed at a room temperature of 22 ± 2 °C and relative humidity of 55 ± 10 % on a light:dark cycle of 12:12 h (artificial light; 5 min twilight transition phase; lights on from 6:00 a.m. to 6:00 p.m.) [14, 31–33]. Pelleted mouse diet (Ssniff rat/mouse maintenance, Spezialdiä ten GmbH, Soest, Germany) and tap water were available ad libitum [14, 31–33]. The animal technician and experimenter were female [14, 31–33]. The mice were handled using a combination of the tunnel and cup method [14, 31–33]. After the experiments, intact male mice could be used for educational purposes and female and castrated male mice were re-homed [14, 31–33].

#### Treatments

This subset comprises four treatments: 1) inhalation anesthesia with isoflurane (age: 10–13 weeks; 26 males, 26 females) [31], 2) injection anesthesia with the combination of ketamine and xylazine (age: 10–13 weeks; 26 males, 22 females) [32], 3) controls/no treatment (age: 10–13 weeks; 13 males, 13 females) [31, 32], and 4) castration (age:18–42 weeks; 19 males, which had previously received treatment 2) or 3) [33]. Further information on the treatments is described in detail in Andresen et al. [14].

#### Image acquisition

Detailed information on the procedure of image acquisition can be found in Hohlbaum et al. [30]. The images were generated in custom-made observation cages (22 cm × 29 cm × 39 cm, three white walls, one transparent wall [14, 31–33]. The floor was covered with 0.5 cm fine wooden fresh and a handful soiled bedding) [14, 31–33]. The mice had access to food pellets and a water bowl placed on the floor of the cage [14, 31–33]. Before images were acquired, the mice were individually transferred into an observation cage where they habituated for 30 min [14, 31–33]. A Canon EOS 350D camera (Canon Inc., Tokyo, Japan) was utilized to take images for approximately 1–2 minutes (in some cases more time was needed) [14, 31–33].

### Subset LW

#### Animals

Female and male mice of the inbred C57BL/6J strain (female: n = 75, male: n = 75) and outbred NMRI strain (female: n = 78, male: n = 78) were utilized for experiments. Mice were either 10 or 48 weeks old (± 5 d) at the start of experiments. Both mouse strains were from in-house breeding in the central laboratory animal husbandry at the Max Rubner Laboratory of the German Institute of Human Nutrition (DIfE), Germany. Breeding, housing and experimental conditions met the specified pathogen free (SPF) conditions. Animal experimentation was approved by the State Office for Occupational Safety, Consumer Protection and Health in Brandenburg (Landesamt für Arbeitsschutz, Verbraucherschutz und Gesundheit – LAVG; permit number: 2347-14-2019). Other data sets from the same study, including 10-week-old female and male C57BL/6J mice, have already been published [34, 35].

#### Housing and husbandry

Females were group housed (n = 5) in conventional type III cages (800 cm^2^, EHRET GmbH Life Science Solutions, Freiburg, Germany) while males were single-housed in open cages of type II (26.8 cm × 21.5 cm × 14.1 cm, Tecniplast, Buguggiate, Italy) due to aggressive behavior against conspecifics. Olfactory as well as visual contact between mice was maintained at all times, because open cage systems were used. Enrichment material in home cages and control cages included aspen wood bedding material (grain size: 2–5 mm, height: 1.5 mm, ssniff Spezialdiäten GmbH, Soest, Germany), 1 cotton nestlet (5 cm x 5 cm, ZOONLAB GmbH, Castrop-Rauxel, Germany), 2 cellulose tissues (green, H3-towel system classic, 23 cm x 24.8 cm, Essity Professional Hygiene Germany GmbH, Mannheim, Germany), 1 aspen wood gnawing bar (100 x 20 x 20 mm, ssniff Spezialdiäten GmbH, Soest, Germany), and 1 cardboard house (16 cm x 12 cm x 8 cm, LBS Biotechnology, United Kingdom). The animal housing conditions were standardized (room temperature: 23 ± 1 °C, relative humidity: 50 ± 10 %, light:dark cycle: 12:12 h of artificial light; lights on: 06:00 a.m. to 06:00 p.m.). Acidified water (pH 2.5–3.0) and autoclaved feed pellets (rat/mouse maintenance, V 1534-300, ssniff Spezialdiä ten GmbH, Soest, Germany) were provided ad libitum. To minimize stress reactions, two animal caretakers and two experimenters handled mice only.

#### Treatments

The animals were assigned into three experimental groups including controls (female: n = 5, male: n = 5), Metabolic Cage Type Number 1 (MC1; female: n = 10, male: n = 10), and Metabolic Cage Type Number 2 (MC2; female: n = 10, male: n = 10). In order to investigate the effect of single-housing, the control mice were kept single in open type II cages. A comparative study was conducted including the investigation of two different metabolic cage types, i.e., MC1 and MC2. For baseline measurements, mice were shortly single-housed in open cages of type II in front of an inserted cage wall (C57BL/6J: white, NMRI: light gray). The other four time points examined were just before expiration of the respective 24 h restraint in different cage systems (control/MC1/MC2). After each 24 h restraint, mice were returned to their home cages with familiar group constellations except for males. After a 6-d resting period, mice were transferred into either control cage, MC1 or MC2 again. Both mouse strains were placed in one of the three cages either twice or four times.

#### Image acquisition

Photographs were taken between 8:00 a.m. and 10:00 a.m. in the setting Shutter Priority (exposure time: 1/1000 s) and without flash adjustment. A camera (*α*6000; Sony, Tokio, Japan) with an interchangeable lens (70 mm, F2.8 DG, MACRO, filter size: 49 mm; Sigma GmbH, Rödermark, Germany) was utilized. Three pictures per mouse were selected for scoring afterwards, ideally displaying the front, left and right side of the mouse face.

### Subset MR

#### Animals

Ninety-six (96) male adult DBA/1JRj mice aged 8–11 weeks, were obtained from Janvier Labs (Saint-Berthevin Cedex, France), kept in the institute’s housing facility (Fraunhofer IZI-MWT, Halle (Saale), Germany). The animals were free of all viral, bacterial, and parasitic pathogens listed in the FELASA recommendations [25]. Animal experimentation was approved by responsible state authority of the state of Saxony-Anhalt (Landesverwaltungsamt Halle; permit number 42502-2-1472).

#### Housing and husbandry

The mice were delivered in large transportation boxes and, in order to avoid the disruption of social structures, subdivided into groups of 4–5 mice, without assembling mice from different boxes. Subsequently, mice were kept in APET IVC type II long cages (36.5 cm x 20.7 cm x 14.0 cm, Tecniplast, Buguggiate, Italy). In case of observed aggression, mice were separated into single housing. Wooden bedding material (spruce granulate, Safe® FS 14, J. Rettenmaier & Söhne GmbH & Co. KG, Rosenberg, Germany), enrichment material (Sizzle Nest, Plexx, Elst, Netherlands) and red plastic igloos (11.4 cm x 5.7 cm, Plexx) were provided. Atmospheric room conditions were standardized at 22 °C ± 2 °C; and 50 ± 20 % air humidity by central air conditioning, light-dark cycles at 12:12 h (06:15 a.m. – 06:15 p.m.) with 30 min twilight transition period. Pelleted mouse diet (Ssniff rat/mouse maintenance, Spezialdiäten GmbH, Soest, Germany) and drinking water quality tap water were available ad libitum. The animal technician and experimenter were female. Mice were handled via their tail and using the cup method. All experiments included final endpoints for further blood analysis.

#### Treatments

Mice received LPS (serotype E.coli O55:B5, Sigma-Aldrich, St. Louis, USA) i.p. in 5 µl volume per g body weight at one of 3 different dosages (250 µg/kg, 1,000 µg/kg, 2,000 µg/kg) or mere 0.9 % phosphate buffered saline (PBS, Gibco, Carlsbad, USA) through a sterile 1 ml syringe (Braun Injekt-F, 26G x 1/2”, B.Braun, Melsungen, Germany), forming 4 experimental groups to test a dose dependency of apparent symptoms, sensitivity of this mouse strain and control for procedural impairment of wellbeing. LPS was stored at a stock concentration at 1 mg/ml and diluted with further PBS at the day of experiment, when mice were weighed and the respective dosage calculated.

#### Image acquisition

Six to eleven photographs were taken per mouse and time point using a Nikon D5100 (Nikon Corp., Tokio, Japan), from lateral left, right and frontal perspective. Therefore, mice were posed under a 10 cm diameter glass dome and habituated for 1 minute, before image acquisition. On the day prior to the experiment, the mice were habituated under the dome for 3 minutes without any interference. Images were taken at baseline (no injection) and 1 h, 2 h, 4 h, 8 h, and 24 h post injection.

### Frame Selection and Face Detection

In subsets based on image data acquired through video recordings, e.g., subsets AW and JW, not every frame of the video was used for MGS evaluation or the pretext task. Since mice usually move around and their faces are not visible in all frames of a video, a fully automatic face detector was applied to ensure that the facial features needed for MGS evaluation were visible. It was published on GitHub under https://github.com/tub-cv-group/mouse-face-detector. The face detector was based on DeepLabCut (DLC) [23] detectors of the eyes, ears, and nose trained on about 400 hand-labeled examples each. A specialized detector was trained for each video dataset. For every recording (i.e., one mouse at a certain time point in the experiment), all frames were ranked according to the number of facial features detected (above likelihood threshold of 0.5). Those with the maximum number of detected features were collected in a list and the sum of the detection confidences for their facial features was computed. Then a minimum time difference of 80 frames between neighboring picks was enforced by removing frames too close to a neighboring one with a higher confidence sum. This time distance ensured that a sufficiently large variety of images could be obtained from each video clip. If the resulting list was shorter than the desired number of frames, the process was continued with a list of frames in which fewer than the maximum number of facial features were detected. They were ranked by detection confidence sum and one by one inserted into the list while enforcing the minimum distance in time. This process was repeated until the desired number of frames was achieved. For the subset AW this was 15 frames, for the subset JW 10 frames per recording.

From the resulting list, the locations of the facial features were used to determine a bounding box around the face of the mouse. First, a minimal bounding box was drawn around all facial features detected with a sufficient likelihood (i.e., we used a threshold of 0.5). This box was then scaled in width and height by a factor of 1.66, which was determined heuristically to include the full face but not too much of the background. If the detector was not able to find the nose and at least an ear, the larger factor of 2.35 was used instead and ensured the inclusion of the missing part of the face in the crop.

Finally, the face crops were ranked according to a computed blurriness measure. The method is based on Fast Fourier Transform [36] and implemented in OpenCV [37]. The least blurry frames were used for MGS scoring.

The KH, subsets LW and MR stem from still images and no frames had to be picked. For the subset KH, the faces were detected with the face detector described above. In the subsets LW and MR, faces were cropped manually in accordance with the original designs of those studies.

### Impairment Label

In many images, it can be assumed that a mouse with impaired well-being is shown, solely based on the time at which the image was taken. Knowledge of the expected impairment can serve as a pretext task for the model training. The impairment label (i.e., “well-being impaired” or “well-being unimpaired”) was assigned, as follows: If the median and the 25th percentile of the human assigned MGS labels at a time point was higher than 0.67, all images from that time point were assigned to the class with “impaired well-being”. In contrast, if the median and the 75th percentile of the MGS labels were lower than 0.67, all images from that time point were assigned to the class with “unimpaired well-being”. All images from time points where neither of these two conditions was met were not assigned any of the impairment labels (i.e., these time points were unclear).

### MGS Label

The MGS by Langford et al. [9] consists of five facial action units (i.e., orbital tightening, nose bulge, cheek bulge, ear position, and whisker change), each evaluated on a 3-point-scale (0 = not present, 1 = moderately visible, 2 = severe).

For images in the five subsets, the average MGS score was calculated across all facial action units. Specifically, a simple average was computed over all valid values (0, 1 or 2), with unrecognizable facial action units excluded and the average taken over all remaining values. An average MGS score was attached to each image, for which at least one human rater assigned values.

Fig 2 illustrates the distribution of human-generated (average) MGS scores for each subset. The distributions of human-generated MGS scores for each facial action unit and subset separated are plotted in Fig 3.

**Fig 2.**
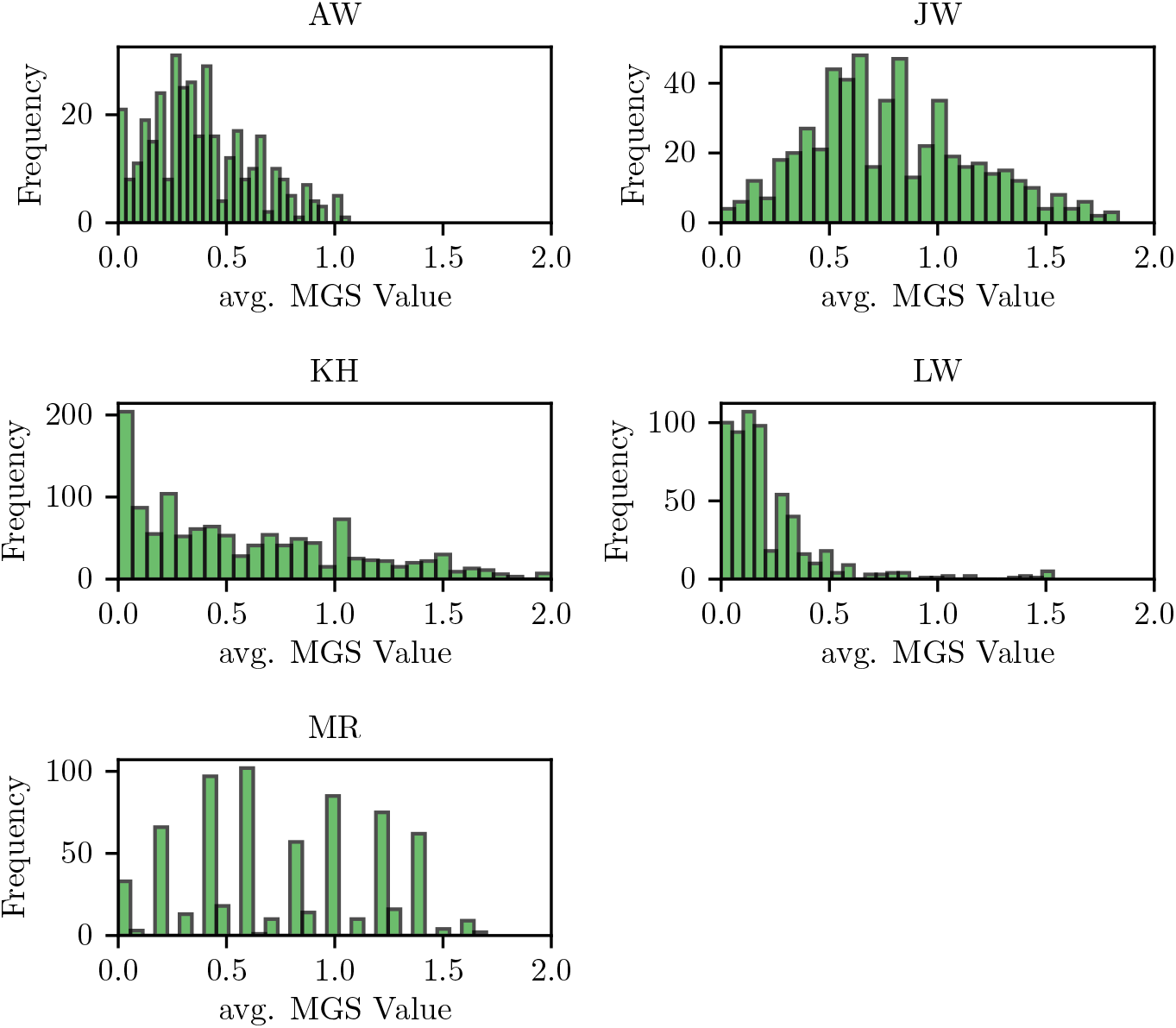
MGS distribution. Histograms showing the distributions of the average Mouse Grimace Scale (avg. MGS) values in the five subsets.

**Fig 3.**
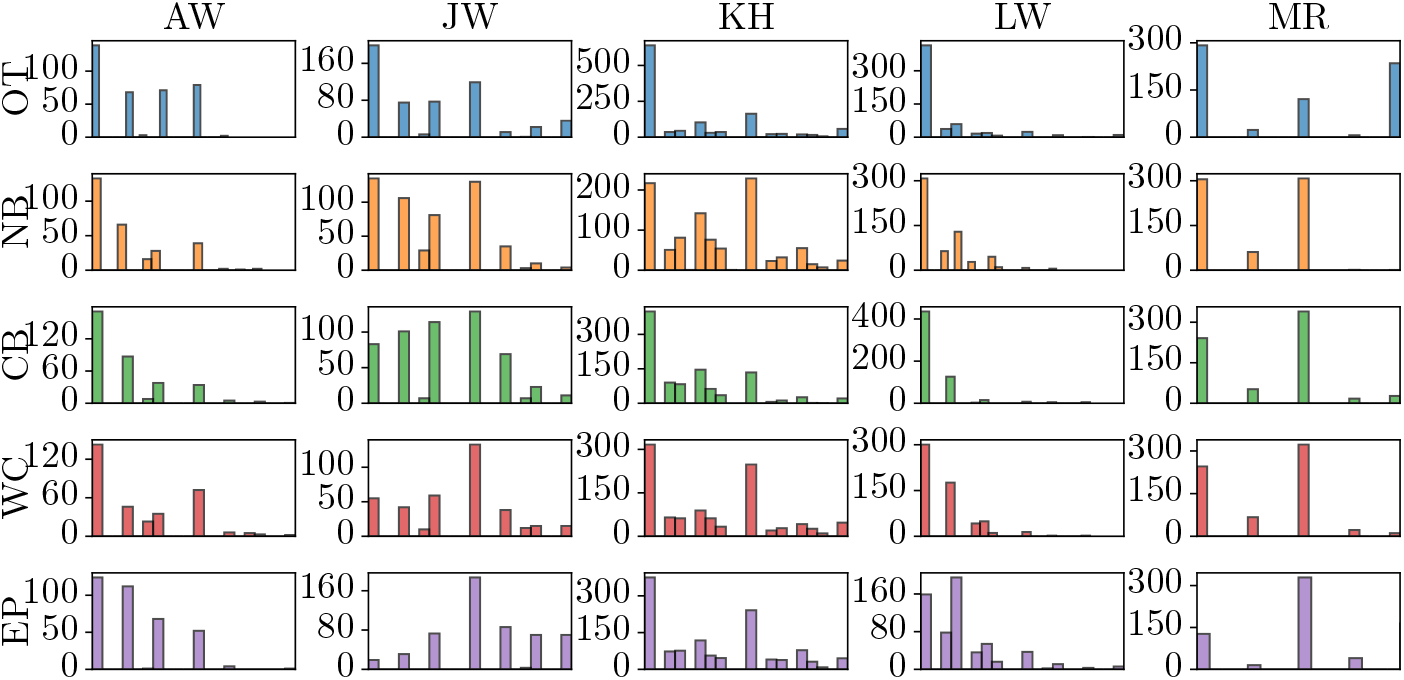
Detailed MGS distribution. Histograms showing the distributions of score of each FAU in the five different subsets AW, JW, KH, LW, and MR. The x-axis of each plot shows the score range from 0 to 2, the y-axis shows the number of images.

### Inter-Rater Reliability

We evaluated the inter-rater reliability (IRR) of the MGS scores for all subsets. A value was computed for each combination of raters that scored the same set of images.

Cohen’s kappa was used for two raters (i.e., a set of images was scored by the same two raters), Fleiss’ kappa for more than two raters. Combinations of raters with less than three examples in common were ignored. Images with only one rater were also ignored for IRR calculation. The IRR values were then averaged with a weight according to the number of images in the group.

For Cohen’s Kappa the implementation in scikit-learn [38] was used. The score is defined as *κ* = (*p*_0_ − *p*_*e*_)*/*(1 − *p*_*e*_), where *p*_0_ is the observed agreement ratio, and *p*_*e*_ is the expected agreement when labels are random. *p*_*e*_ is estimated with a per-annotator empirical prior over the class labels. For Fleiss’ Kappa [39] the implementation in statsmodels [40] is used. The score is computed as 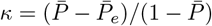 where 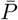 is the average observed proportion of agreement over all pairs of raters and 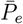 denotes the expected proportion of agreement in the case of random labels.

For comparison, the intra-class correlation coefficients (ICC) were also computed because these values are often reported in related literature. However, none of the cases and sets of assumptions made for ICC to be applicable is fitting to this kind of data. The set of raters in most of the subsets used in the present work was neither fixed nor uniformly random since the MGS-labeled subsets were obtained from different laboratories. Therefore, the two-way ICC^1^ could not be applied to whole datasets.

Instead, it was computed for each occurring number of raters. I.e., for each image the number of raters was counted and images with the same number of raters were grouped. The group of images with only one score was not taken into account. For each group, the ICC was calculated and then a weighted average was computed over all acquired ICC values. The weight was the number of images on which the value was based. This was performed for each facial action unit separately.

### Neural Network Model

#### Architecture

The proposed backbone is the ResNet 50 model [41]. It is an established standard model for deep neural networks. The optimization approach is based on transfer learning. The final model for the main task is achieved in several steps, in each of which the weights are optimized for a different task, with the goal of achieving a favorable initialization for the next step. Transfer learning helps to learn from smaller datasets, when some aspects of the data could already be learned from a proximate task for which more data is available [42].

The first pretext task is the common object recognition on the ImageNet-21k dataset [43], which contains about 40 million images divided into about 21 thousand classes. The checkpoint is freely available under the name Big Transfer BiT-M-R50x1 and achieved 78.4 % accuracy on the ImageNet test set [44]. The second task involves binary classification between pictures of mice assigned the labels “unimpaired well-being” or “impaired well-being”. The latter are pictures in which the facial expression is expected to show increased MGS scores. The labels were created as described in the section on the “impairment label”.

The final task was to predict the average MGS value. Since the five facial action units are each scored with 0, 1 or 2, the average was a floating point number between 0 and 2. Most images with MGS scores were evaluated by several human raters and the simple average of all available scores was taken.

#### Network Heads

The base model is a ResNet 50 [41] initialized with ImageNet 1k V1 weights. For the binary classification task a multi-layer perceptron is replacing the output layer. It consists of dropout, a linear layer with 128 neurons and ReLU activation and the output layer with one neuron and sigmoid activation. The head is smaller for the regression task, when it only consists of dropout and an output neuron with linear activation.

#### Training

Whenever the head of the neural network model was exchanged for a new one for the next task, the new zero initialized weights were first trained to convergence before the rest of the network was unfrozen. This procedure prevents the large gradients due to the random initialization to destroy the pre-trained state of the backbone. Input images were randomly augmented through a variety of techniques using RandAugment [45]. In the final configuration, we trained all models for 200 epochs, which ensures convergence.

## Results and Discussion

Most performance or error estimates were computed based on a held-out test set consisting of the images of 10 % of the animals drawn at random (Table 3). For the cross-dataset evaluation, the test set consisted of full subsets not used for training (Table 4, Table 5). The MGS regression results presented are grouped into these two cases and given as root mean squared error (RMSE), inter-rater RMSE, and Pearson Correlation Coefficient.

**Table 3.**
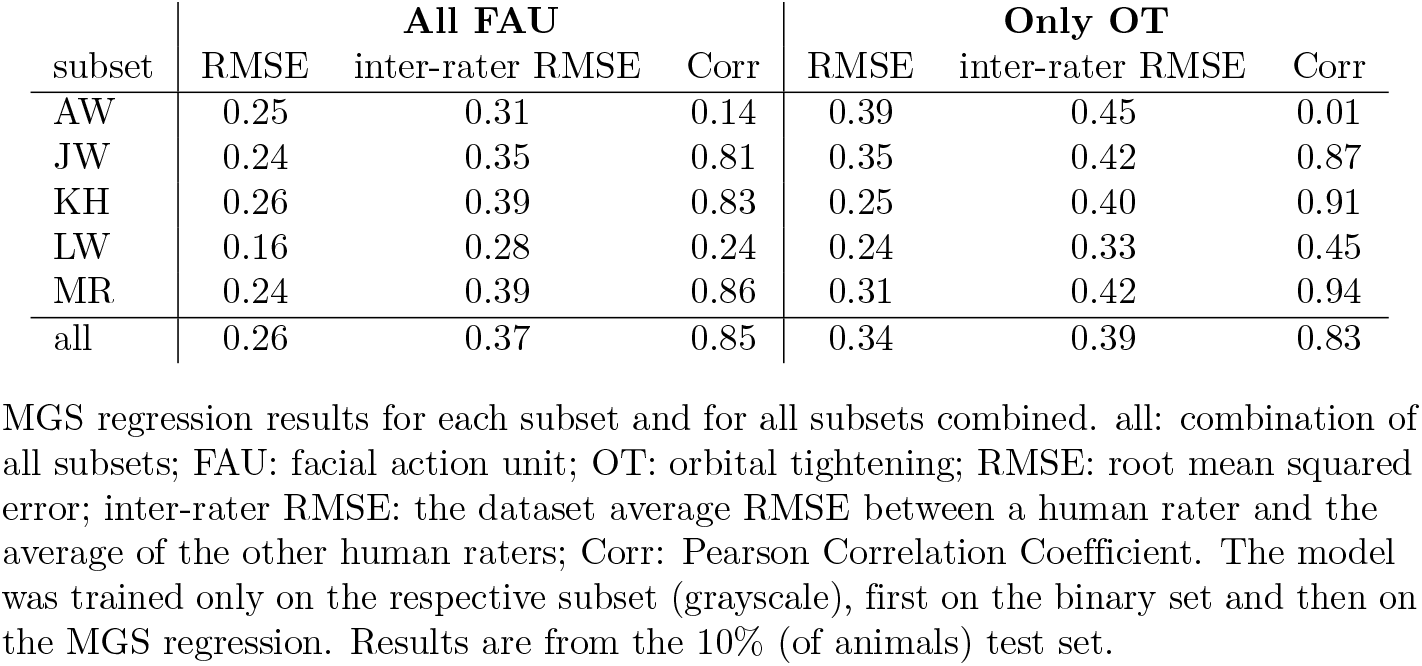
MGS regression results.

| subset | All FAU |  |  | Only OT |  |  |
| --- | --- | --- | --- | --- | --- | --- |
|  | RMSE | inter-rater RMSE | Corr | RMSE | inter-rater RMSE | Corr |
| AW | 0.25 | 0.31 | 0.14 | 0.39 | 0.45 | 0.01 |
| JW | 0.24 | 0.35 | 0.81 | 0.35 | 0.42 | 0.87 |
| KH | 0.26 | 0.39 | 0.83 | 0.25 | 0.40 | 0.91 |
| LW | 0.16 | 0.28 | 0.24 | 0.24 | 0.33 | 0.45 |
| MR | 0.24 | 0.39 | 0.86 | 0.31 | 0.42 | 0.94 |
| all | 0.26 | 0.37 | 0.85 | 0.34 | 0.39 | 0.83 |

MGS regression results for each subset and for all subsets combined. all: combination of all subsets; FAU: facial action unit; OT: orbital tightening; RMSE: root mean squared error; inter-rater RMSE: the dataset average RMSE between a human rater and the average of the other human raters; Corr: Pearson Correlation Coefficient. The model was trained only on the respective subset (grayscale), first on the binary set and then on the MGS regression. Results are from the 10% (of animals) test set.

**Table 4.** RMSE results for the transfer to other subsets.

| RMSE | All FAU |  |  |  |  | Only OT |  |  |  |  |
| --- | --- | --- | --- | --- | --- | --- | --- | --- | --- | --- |
|  | AW | JW | KH | LW | MR | AW | JW | KH | LW | MR |
| AW |  | 0.59 | 0.50 | 0.30 | 0.58 |  | 0.63 | 0.59 | 0.45 | 0.99 |
| JW | 0.43 |  | 0.59 | 0.62 | 0.50 | 0.57 |  | 0.71 | 0.72 | 0.91 |
| KH | 0.50 | 0.42 |  | 0.39 | 0.38 | 0.41 | 0.75 |  | 0.40 | 0.96 |
| LW | 0.27 | 0.63 | 0.57 |  | 0.55 | 0.50 | 0.67 | 0.67 |  | 0.76 |
| MR | 0.37 | 0.40 | 0.43 | 0.49 |  | 0.71 | 0.62 | 0.72 | 0.56 |  |
| all | 0.33 | 0.60 | 0.42 | 0.42 | 0.34 | 0.42 | 0.62 | 0.49 | 0.43 | 0.66 |

MGS regression results for the transfer to other subsets. all: combination of all subsets; FAU: facial action unit; OT: orbital tightening; RMSE: root mean squared error. The model was trained only on the subset in each row (grayscale), first on the binary set and then on the MGS regression. Results are from the whole dataset in the column. The last row indicates results from models trained on all subsets together except the one tested on in the column.

**Table 5.**
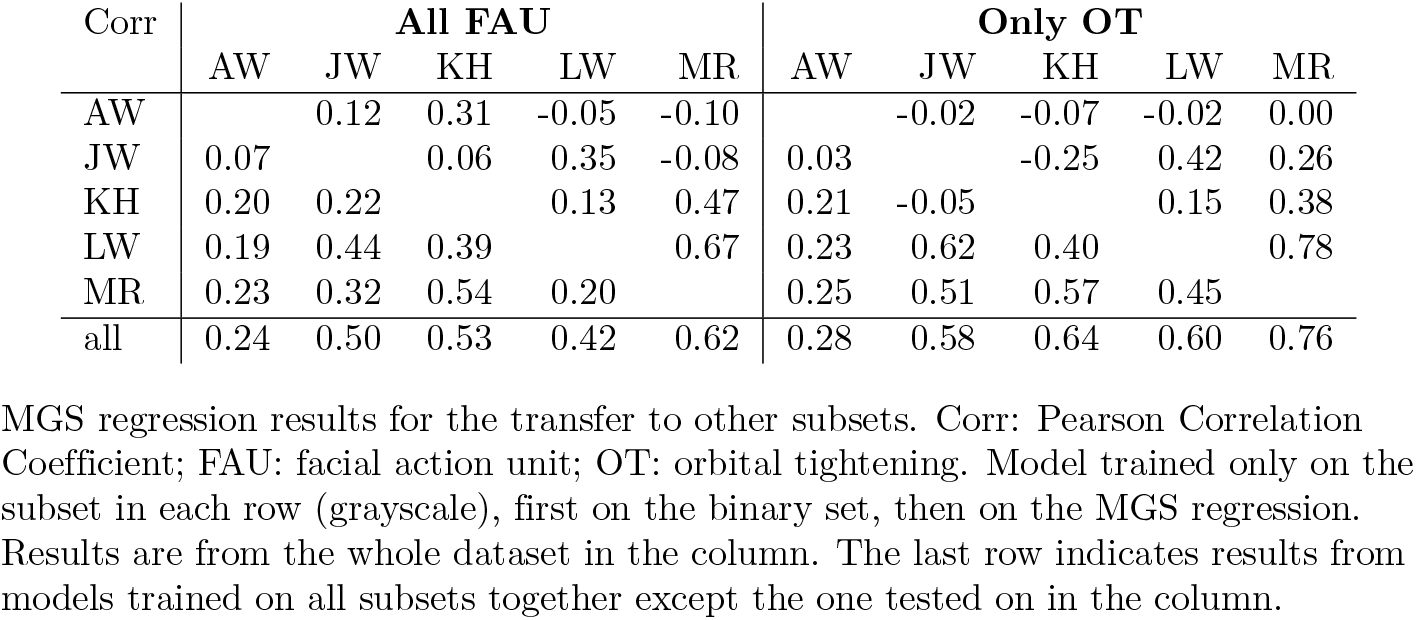
Correlation results for the transfer to other subsets.

A RMSE value can be understood as an average distance on the MGS scale ranging from 0 to 2 between the human-given score and the model output. The best possible RMSE is 0 and the higher it is, the worse the model performs. Because of the square operation, the RMSE is sensitive to large outlier errors. The inter-rater RMSE is a statistic about the human-given scores. It is computed by leaving out one human expert’s score and computing an average over the remaining raters. Then a RMSE difference between the two values is calculated giving an average difference between any human rater and the mean of the other raters, i.e., a measure of how one expert differs from his or her colleagues. It serves as a lower bar for assessing the RMSE values of the neural network model, which may not predict the exact MGS values for all images but is supposed to be at least as close as human experts would be. The Pearson Correlation Coefficient for the correlation between model output and human-given scores is another measure of the model’s performance. The best value is 1 and the worst would be -1.The correlation is especially useful when comparing two different models trained on two subsets, which have very different score distributions. Here the RMSE will always favor the model trained on a narrower score distribution, because just predicting close to the mean will lead to a low RMSE, while being independent of image contents. The correlation coefficient is per definition normalized and thus captures the connection between score and model output independent of the variance in the dataset. It is, however, blind to a systematic bias, which the RMSE would show.

### Rater Agreement

The rater agreement varied based on subsets and facial action units (Table 2). The IRR was highest in the subset MR for orbital tightening (0.73) and lowest in the same subset for whisker change (-0.07). Averaging over the five facial action units yielded the best IRR in the subset JW (0.42), followed by the subset AW (0.33), and the worst in the subset LW (0.13). Orbital tightening has the highest reliability score in all subsets, thus being the easiest for raters to judge in agreement with each other. The other facial action units generally exhibited fair agreement in some subsets and only slight agreement in others. The ICC mostly showed the similar qualitative results as the IRR with some exceptions.

The differences between the subsets found in the IRR/ICC values can be explained by the different approaches to train the human raters. MGS training at the highest intensity was performed for the subsets AW and JW, potentially contributing to their higher average IRR and ICC values. Briefly, the training consisted of three group training (approximately 1 to 2.5 hours) and two individual training sessions. In the first group training session, the trainer explained the MGS to the trainees using the original MGS manual by Langford et al. [9], the MGS poster provided by the NC3Rs (National Centre of Replacement, Refinement, and Reduction, United Kingdom of Great Britain: https://nc3rs.org.uk/3rs-resource-library/grimace-scales/grimace-scale-mouse), and a MGS poster including images from the subsets AW and JW. In the individual training sessions, trainees were asked to score approximately 40 images from the subsets AW and JW. Discrepancies between the MGS scores generated by the trainees were addressed during the group training sessions, where trainees explained to one another the reasons behind their decisions and reached a consensus. The sessions also included instructions on identifying images that should not be scored using the MGS, as certain behaviors can influence the mice’s facial expression.

### MGS regression for each subset and the combination of all subsets

Depending on the subset, the RMSE ranged from 0.16 (LW) to 0.26 (KH) on the MGS scale with values from 0 to 2. This means that the model, trained and tested on the subset LW, deviated on average by 0.16 from the human-labeled MGS scores. The RMSE of models trained and tested on the other subsets KH (0.26), AW (0.25), JW (0.24), MR (0.24), or all subsets (0.26) was slightly higher compared to LW. A reason for this observation may be the difference in the distribution of the average MGS scores across the five subsets (Fig 2), with the subset LW consisting mainly of images with low MGS values (*<* 0.5) and only a few images with higher MGS values. In contrast, the other subsets exhibit a wider distribution of MGS scores than the subset LW. If a dataset contains only images with low average MGS scores, and the distribution of these scores is narrow, this may lead to a low-score bias, which in turn makes it easy to achieve a lower average error (RMSE). Sturman et al. also reported the issue of low-score bias for their dataset [21].

To better interpret the RMSE between the model and the human raters, the value can be compared with the inter-rater RMSE. In Table 3, the average RMSE of the subsets between a human rater and the average of the other human raters is demonstrated. The RSME from the respective other raters varies between 0.28 (LW) and 0.39 (MR). With this magnitude of errors, the model is closer to an average human rater than a human rater is from his or her colleagues.

For a more comprehensive evaluation, it is also useful to consider the Pearson Correlation Coefficient between the prediction of the model and the average human-generated MGS score (Table 3). The values range between 0.14 (AW) and 0.86 (MR). Although the subset AW demonstrated a very weak correlation between the model’s prediction and the human-generated MGS scores, the RMSE value did not indicate such an outlier performance. As discussed above, this may be also due to the narrow distribution of MGS scores in this subset. With MGS scores ranging from 0 to 1.25, it exhibits the second-lowest distribution following subset LW, for which the Pearson Correlation Coefficient is also weak (0.24). In an extreme case, the model could consistently predict a value close to the average score and still achieve a relatively low RMSE, despite having no correlation with the human-generated MGS score. The Pearson Correlation Coefficient of the other subsets, including a wider distribution of MGS scores, indicated a very strong positive correlation (*>* 0.8). The same was true for the model trained on the combination of all subsets. Similar results were obtained by Sturman et al., who reported a Pearson Correlation Coefficient of 0.87 [21].

### Cross-dataset evaluation

To determine whether models trained on one or more subsets could be used to predict MGS scores of another subset that was not included in the training, we carried out a cross-dataset evaluation. In general, this approach yielded higher errors compared to models trained and tested on parts of the same subset(s) (Table 4). The models trained on the subsets KH (RMSE: 0.38–0.50) or MR (RMSE: 0.37–0.49) showed the best performance when transferred to the other subsets. As the subset KH included mice with black fur and the subset MR mice with dilute-brown, the generalization seemed to perform well when mice with these fur colors were included in the training dataset. It should be noted that all images were converted to grayscale. The treatments administered to the mice in the subsets KH and MR may also be influential for generalization, as the treatments induce different pharmacological effects or affective states, which can affect facial expression to varying extents. The influence of the five facial action units differs across various states, such as illness and pain [9, 46]. This may result in some facial action units being assigned lower and others higher MGS scores, depending on the treatment. In a previous study, it was assumed that the treatments can result in a different facial expression [14]. Both subsets KH and MR seemed to represent a wide range of MGS scores for each facial action unit (Fig 3). However, this was also found for the model trained on the subset JW, which exhibited poor transferability to the other subsets, probably due to pronounced differences from the remaining subsets, such as the white fur coloration and/or the use of wound clips to close the incision site.

Although the transfer of models trained on the subsets AW or LW resulted in high RMSE values up to 0.59 or 0.63, there was one exception: interestingly, models trained on the subsets LW or AW and tested on the respective other subset achieved the lowest RMSE values for the cross-dataset evaluation, i.e. 0.27 for the subset LW and 0.30 for the subset AW. This discrepancy may be attributed to the narrow distribution of MGS scores in these two subsets, as discussed above, in contrast to the wider distribution observed in the other subsets. If a model is trained exclusively on low MGS scores, it will not generalize well to predict higher MGS scores. The highest RMSE values were found for a model trained on LW and tested on JW, and vise versa. A part of this is likely due to the mentioned lack of high scores in the subset LW, which leads LW-trained models to be insensitive to high scores. Another factor is the generally low reliability of the MGS scores in the subset LW (Table 2).

The combination of several subsets for training improved the generalization and resulted in RMSE values ranging between 0.33 and 0.60 (Table 4). Considering the Pearson Correlation Coefficient alongside the RMSE, the model trained on all subsets except the subset MR and tested on the latter demonstrated the best overall performance. However, it is difficult to identify the underlying reasons for this result. A notable difference compared with the other subsets was the smaller size of the observation cage in the recording setup (i.e., 10 cm in diameter), along with the use of a uniform background, which may partially account for the enhanced generalization performance.

Overall, the Pearson Correlation Coefficient revealed a weaker correlation between the model’s prediction and the human-generated MGS scores for the cross-dataset evaluation (Table 5) when compared to models trained and tested on the same subset (Table 3). This means that the transfer to other subsets works inconsistently. Combining multiple subsets into a larger training set provided some degree of stability but did not achieve the performance level of models trained and tested on the same subset.

### Focusing on orbital tightening

Considering that orbital tightening achieved the highest IRR in all subsets (Table 2), we hypothesized that the models would perform better when exclusively orbital tightening is used for training and has to be predicted. Therefore, we further adopted an approach that concentrated solely on this facial action unit (Table 3, Table 4, Table 5). However, contrary to our expectations, RMSE values indicated a higher average distance on the MGS scale between the human-given scores and the model output when compared to the approach including all facial action units (Table 3). This observation was consistent across most models, regardless of whether they were trained on a single subset or multiple subsets (Table 3), or transferred to another subset (Table 4). The RMSE was slightly better using only orbital tightening for the subset KH, otherwise it was worse (Table 3). The correlation, however, increased for all subsets except AW (Table 3). In the cross-dataset evaluation, a similar trend was observed (Table 5). This situation with a comparatively higher RMSE and higher correlation using only orbital tightening might indicate a better fit but also the existence of a bias.

Sturman et al. reached a correlation of 0.8047 between model prediction and human-generated score for orbital tightening in black-furred mice [21], which is comparable to the results of the present study. When all subsets were included in the training of the (only OT) model, a correlation of 0.83 was achieved (Table 3). Another study using black-furred mice reported for a similar task a mean absolute error (MAE) of 0.871 using a fine-grained scale from 0 to 9 for orbital tightening [15]. Converted to a scale of 0 to 2, this would correspond to a MAE of 0.194. With the (only OT) model presented here, a similar MAE of 0.24 was achieved.

## Conclusion

This study introduces a diverse dataset containing approximately 35,000 images of mouse faces, with more than 3,000 images labeled according to the MGS. Moreover, a deep learning model for automated MGS scoring is presented, accompanied by benchmark results.

The convolutional neural network model outputs a MGS floating point value given an image of a mouse face. The output value has an expected error of 0.26 on the MGS ranging from 0 to 2 when the model was trained on all subsets of the dataset. This is a smaller error than the average human rater makes. For effective training of the model, it was essential that the MGS scores of the training data covered the full range from 0 to 2 across all facial action units. Restricting a model to the most reliable facial action unit, OT, did not improve its performance.

When the model was applied to another subset of images, which differed in terms of mouse strain, treatment, laboratory, and/or image acquisition setup from the subset used for training, the performance deteriorated. The error can be reduced when training was conducted using a combination of several subsets of images. Here the variety of irrelevant visual information seems to be ignored more consistently and the model focuses on a more generally applicable definition of a pain face.

For predicting MGS scores for a novel dataset, the most reliable model is the one trained on the combined subsets. However, the best performance is achieved when human raters generate MGS scores for a subset of the novel dataset to fine-tune the existing model. Future research can demonstrate improvements over the present results in a readily comparable manner by using the dataset published alongside this study.

## Acknowledgments

Funded by the Deutsche Forschungsgemeinschaft (DFG, German Research Foundation) under Germany’s Excellence Strategy – EXC 2002/1 “Science of Intelligence” – project number 390523135. We thank Ines Koska for assisting in the study of the subset MR. The authors have no conflicts of interest to declare.

## Footnotes

1 ICC(3,1): two-way mixed

